# A Gridded Microclimate Dataset from a Sub-Arctic Biodiversity Hotspot in Finland

**DOI:** 10.1101/2024.03.30.587419

**Authors:** P. Niittynen, H. Salminen, P. Peña-Aguilera, J. Aalto, J. Alahuhta, M. Luoto, T. Maliniemi, T. Rissanen, T. Roslin, H. Tukiainen, V. Tyystjärvi, J. Kemppinen

## Abstract

Microclimate measurements are crucial for explaining and predicting functions and patterns in nature, yet the availability of microclimate data often poses a challenge. This study introduces a dataset comprising 63 spatially continuous microclimate surfaces for the Kilpisjärvi region in northwestern Finland. The study region is a biodiversity hotspot for arctic-alpine flora and fauna and one of the most extensively investigated regions in Northern Europe. The data were gathered through a collaborative network of microclimate loggers, encompassing 430 measurement locations that comprehensively cover the 300 km2 landscape under study. We employed predominantly well-performing Random Forest models to project microclimate variables across the study area at a 3-metre spatial resolution. The high-resolution and extensive spatial coverage of the dataset facilitates examination of microclimate characteristics and variability across this sub-Arctic region, providing a valuable resource for both theoretical and applied research, as well as for biodiversity conservation.

## 1. Introduction

Microclimate refers to the distinct climatic conditions that impact organisms and ecosystem processes at fine spatiotemporal scales, typically near the ground surface (Chen et al., 1999; Geiger, 1965). Microclimate is mainly regulated by factors such as topography and vegetation, which cause buffered microclimate variability (Bramer et al., 2018; De Frenne et al., 2021). Under specific conditions, such as a thick snowpack, microclimate can even become decoupled from atmospheric conditions (Pepin and Norris, 2005; Zhang, 2005), revealing relationships in nature that would be obscured by more coarse-resolution climate data (Niittynen et al., 2020; Potter et al., 2013). Compared to macroclimate, microclimate better represents the conditions to which organisms are truly exposed, exhibiting pronounced local-scale variability with clear implications for biodiversity research, especially when studying the local effects of climate change (Aalto et al., 2022; Hannah et al., 2014).

Microclimate is a multifaceted phenomenon: different seasons are often characterised by distinct microclimate patterns (Bokhorst et al., 2009; Niittynen et al., 2020); soil and air temperatures can be locally decoupled (Kropp et al., 2016); and mean and extreme temperatures may exert contrasting effects on biotic communities (Bokhorst et al., 2015; Parmesan et al., 2000). Another crucial aspect of microclimate is soil moisture, which also displays great local-scale variability due to the strong topographic control over top-soil moisture (Burt and Butcher, 1985; Kemppinen et al., 2023; le Roux et al., 2013). Arctic and alpine regions, characterised by overall cold temperatures, strong seasonality, and complex terrains, may differ from other ecosystems in regard to the main drivers of microclimate variability (Aalto et al., 2022; Opedal et al., 2015). The absence of a tree canopy means that local radiative conditions and heat transfer are primarily regulated by other factors like slope orientation and snow conditions (Rixen et al., 2022; Winkler et al., 2016).

The Kilpisjärvi region in Northern Finland is marked by large environmental gradients and geodiversity. Known for its high arctic-alpine biodiversity, the region is considered one of the biodiversity hotspots in the Scandic mountains (Kauhanen, 2013). The Kilpisjärvi village has an active research station owned by the University of Helsinki, where decades of intensive study have covered topics such as plant and animal ecology (Aho and Kalela, 1966; Kemppinen et al., 2021a; Peña-Aguilera et al., 2023), geomorphology (Aalto and Luoto, 2014), geodiversity (Salminen et al., 2023), geophysics of ice and snow (Leppäranta et al., 2016; Raidla et al., 2015), and paleolimnology (Kauppila and Salonen, 1997). However, research on the microclimate of the region has gained prominence only recently (Kemppinen et al., 2021b; Tyystjarvi et al., 2022), due to technological advances in affordable logger technology (Maclean et al., 2021; Wild et al., 2019).

This study provides the first comprehensive analyses and ready-to-use dataset covering multiple aspects of microclimate for general use in the Kilpisjärvi region. We utilised microclimate data from 430 study sites spread across the area, collected between 2019 and 2023. From the raw time series of soil temperature, near-surface air temperature, and soil moisture, we constructed 63 microclimate variables. These variables were then related to a broad array of topographical and remotely sensed predictors using machine learning models to produce spatial microclimate layers for the entire study area.

## 2. Material and Methods

All data processing and analyses were conducted in R software (version 4.3.0; R Core Team, 2023).

### 2.1. Study area

The study area consists of 296 km^2^ of sub-Arctic tundra, wetlands and sparse mountain birch forests in northwestern Finland (20.55 – 21.22°E, 68.87 – 69.12°N; **Figure 1**). The elevation varies between 458 and 1029 m a.s.l. within the study area. In addition, variation in local topography, together with heterogeneous snow and moisture conditions result in a mosaic of different vegetation types. The annual mean air temperature is −1.4°C and the total annual precipitation is 516 mm (1991–2020) as measured by the nearby meteorological station of Enontekiö Kilpisjärvi kyläkeskus (Jokinen et al., 2021). The permafrost is discontinuous in the region and occurs mainly deep in the higher mountains. Thus, it has little effect on the near-surface conditions (King and Seppälä, 1987; Westermann et al., 2015).

**Figure 1.**
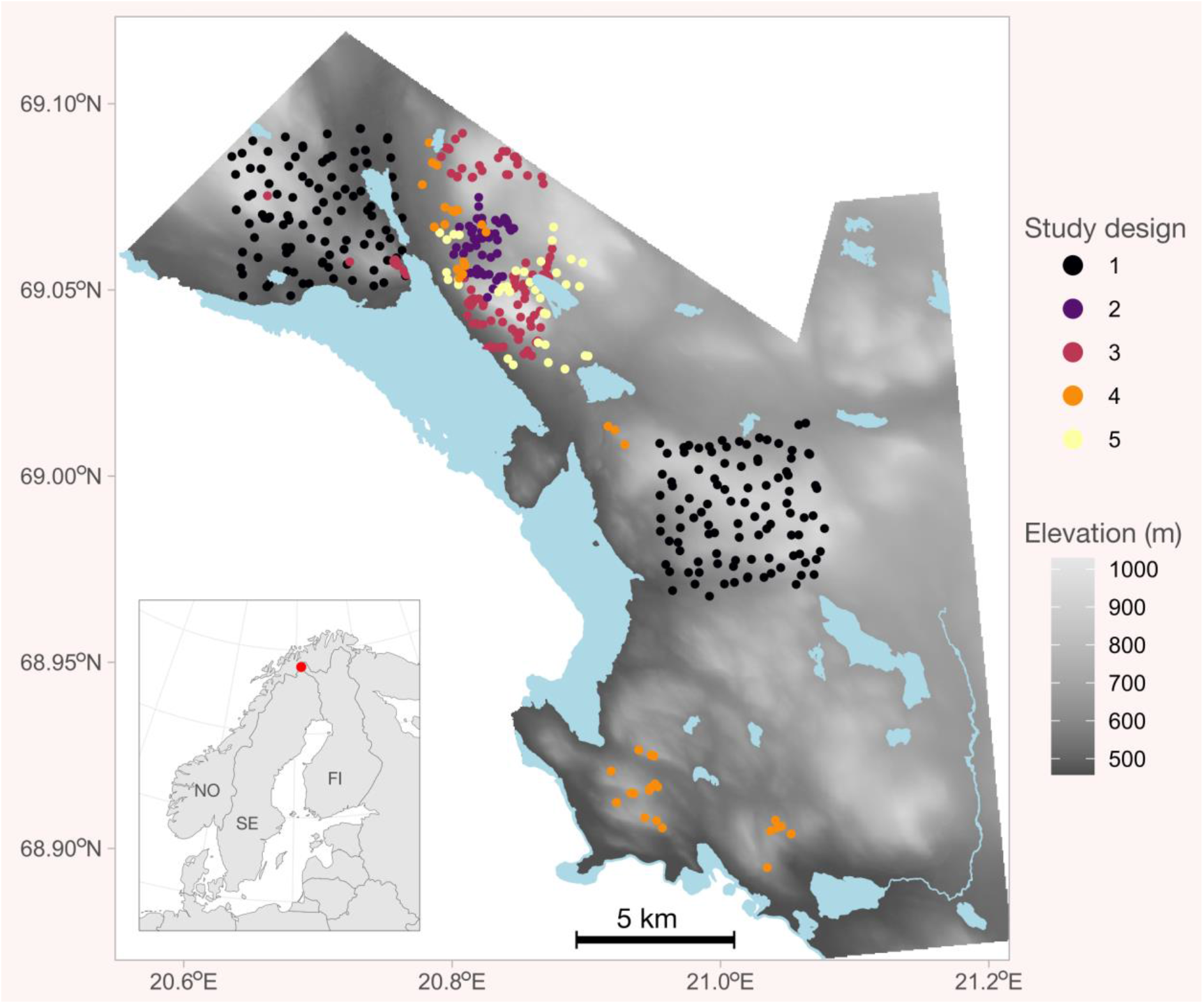
Elevation of the study area and location of the study sites. The colour and numbering of sites refer to the study designs listed in the main text. Light blue areas represent water bodies.

### 2.2. Microclimatic data and variables

All microclimate data are collected using TMS-4 microclimate loggers (Standard datalogger, TOMST s.r.o., Prague, Czechia, Wild et al., 2019). TMS-4 loggers have three temperature and one soil moisture sensor. Here, we utilised the T1 (soil temperature at 6 cm depth), T3 (near-surface air temperature 15 cm above the soil surface) and soil moisture sensors (top 15 cm of the soil). The T2 data (at the soil surface) were not utilised, because they often do not represent conditions comparable across loggers: some loggers were pushed into the soil or moss layer and thus do not represent the surface conditions. Here we included measurements made every 30 minutes.

All logger data were quality-filtered and preprocessed (see detailed descriptions of the data cleaning methods in Aalto et al. 2022 and Kemppinen et al. 2023). In summary, 1) we removed all data outside the period during which the logger was in field; 2) we removed all unrealistically high or low temperatures and moisture values; 3) we cross-correlated the time series across loggers aiming to identify unrealistically high sudden peaks (or reversed drops) in the data and removed them; 4) we plotted all individual time series, one site and month at the time, and checked visually whether there were any clearly erroneous data and removed them; 5) if there were data for a logger from a time when the logger was under stable ‘office’ conditions outside of the field, we used that data to check and correct for potential systematic differences across the three temperature sensors within the logger.

The dataset consists of a total of 430 study sites with loggers spread across the study area (**Figures 1 & S2**). The locations of the majority of the study sites were positioned at centimetre-accuracy with high-accuracy GPS devices and the rest with hand-held Garmin GPS devices at approximately 3 m accuracy. The study sites are part of multiple study designs:

1. 200 loggers were installed around Malla and Ailakkavaara mountains in June 2019. The locations were chosen by stratified random sampling to capture the variation of the main environmental gradients.
2. A set of 50 loggers were installed in June 2018 between mount Saana and Korkea-Jehkas. The locations of the loggers were based on previous soil moisture investigations in a systematic study design (Kemppinen et al., 2018), with attention to cover the entire soil moisture gradient. The study design was further complemented by adding sites to places with extreme soil moisture and snow conditions, and to higher elevations near mountain tops.
3. A set of 105 loggers were installed in summer 2020, on the slopes of Saana, Malla and Jehkas mountains. These locations were designed to complement the two above-mentioned study designs and mainly consist of subjectively selected locations with a presence of rare and endangered plant species.
4. In August-September 2020, a set of 42 loggers were installed across several mountains. Logger locations were based on an approximate location of an old vegetation plot data (Haapasaari, 1988) that was relocated for several research purposes (Maliniemi et al., 2018; Salminen et al., 2023) and that is being monitored for temporal plant community changes. Loggers are located in dry dwarf shrub tundra heaths and cover mesotopographically differentiated locations.
5. In June 2020, 35 loggers were deployed in the area between Lake Kilpisjärvi and Mount Jehkas. The study aimed to investigate how plant and arthropod communities are shaped by microclimatic conditions. A thorough stratified random design was implemented, encompassing a wide range of environmental conditions in terms of topography, vegetation height, snow depth, and distance to water bodies.

Altogether, we calculated 63 microclimatic variables from the logger data (**Table 1**). All variables are based on data from 2019-09-01 to 2023-08-31, thus covering four years (Figure 2). While calculating the microclimatic variables, we divided the data first into the four hydrological years (i.e., September - August), calculated the variables for each year separately, and finally, averaged the variables over the four years. Soil moisture was considered only during the summer months (June - August) and only during days when the soil temperature was above 1℃.

**Table 1.** List of microclimatic variables calculated from the daily mean, minimum and maximum microclimate time series. T1 = soil temperature. T3 = near-surface air temperature. Moist = soil moisture. VWC = volumetric water content.

| Name | Sensor | Explanation | Unit |
| --- | --- | --- | --- |
| T1_mean | T1 | Mean annual soil temperature | °C |
| T3_mean | T3 | Mean annual near-surface air temperature | °C |
| T1_mean_1 ...<br>T1_mean_12 | T1 | Mean monthly soil temperatures | °C |
| T3_mean_1 ...<br>T3_mean_12 | T3 | Mean monthly air temperatures | °C |
| T1_min | T1 | Minimum soil temperature (0.5 percentile of daily minimum temperatures) | °C |
| T1_max | T1 | Maximum soil temperature (99.5 percentile of daily maximum temperatures) | °C |
| T3_min | T3 | Minimum air temperature (0.5 percentile of daily minimum temperatures) | °C |
| T3_max | T3 | Maximum air temperature (99.5 percentile of daily maximum temperatures) | °C |
| T1_TDD | T1 | Thawing degree days, i.e., the sum of daily mean soil temperatures when mean soil temperature > 0°C | °C days |
| T3_TDD | T3 | Thawing degree days, i.e., the sum of daily mean air temperatures when mean air temperature > 0°C | °C days |
| T1_GDD3 | T1 | Growing degree days, i.e., the sum of daily mean soil temperatures when mean temperature > 3°C | °C days |
| T3_GDD3 | T3 | Growing degree days, i.e., the sum of daily mean air temperatures when mean temperature > 3°C | °C days |
| T1_GDD5 | T1 | Growing degree days, i.e., the sum of daily mean soil temperatures when mean temperature > 5°C | °C days |
| T3_GDD5 | T3 | Growing degree days, i.e., the sum of daily mean air temperatures when mean temperature > 5°C | °C days |
| T1_FDD | T1 | Freezing degree days, i.e., the sum of daily mean soil temperatures when mean temperature < 0°C | °C days |
| T3_FDD | T3 | Freezing degree days, i.e., the sum of daily mean air temperatures when mean temperature < 0°C | °C days |
| T1_TDD_days | T1 | Thawing degree days count, i.e., the number of days when mean soil temperature > 0°C | days |
| T3_TDD_days | T3 | Thawing degree days count, i.e., the number of days when mean air temperature > 0°C | days |
| T1_GDD3_days | T1 | Growing degree days count, i.e., the number of days when mean soil temperature > 3°C | days |
| T3_GDD3_days | T3 | Growing degree days count, i.e., the number of days when mean air temperature > 3°C | days |
| T1_GDD5_days | T1 | Growing degree days count, i.e., the number of days when mean soil temperature > 5°C | days |
| T3_GDD5_days | T3 | Growing degree days count, i.e., the number of days when mean air temperature > 5°C | days |
| T1_FDD_days | T1 | Freezing degree days count, i.e., the number of days when mean soil temperature < 0°C | days |
| T3_FDD_days | T3 | Freezing degree days count, i.e., the number of days when mean air temperature < 0°C | days |
| T3_frost | T3 | The sum of daily minimum air temperatures when daily minimum temperature < 0°C between May 1st and August 15th and when mean soil temperature > 1°C (i.e., growing season) | °C |
| sos1 | T1 & T3 | Start of season as the first day of year when both mean soil and air temperature are over 1°C | Day of year |
| sos3 | T1 & T3 | Start of season as the first day of year when both mean soil and air temperature are over 3°C | Day of year |
| eos1 | T1 & T3 | End of season as the last day of the year when both mean soil and air temperature are over 1°C | Day of year |
| eos3 | T1 & T3 | End of season as the last day of the year when both mean soil and air temperature are over 3°C | Day of year |
| T1_ft | T1 | Freeze-thaw cycles as the number of times when soil temperature crosses 0°C and the amplitude of the temperature change is > 0.5°C. Calculated from daily minimum and maximum temperatures. | Count of events |
| T3_ft | T3 | Freeze-thaw cycles as the number of times when air temperature crosses 0°C and the amplitude of the temperature change is > 1°C. Calculated from daily minimum and maximum temperatures. | Count of events |
| moist_mean_summer | Moist | Mean summertime soil moisture. Calculated over days when mean soil temperature is > 1°C in June-August | VWC% |
| moist_max_summer | Moist | Maximum summertime soil moisture. Calculated over days when mean soil temperature is > 1°C in June-August | VWC% |
| moist_min_summer | Moist | Minimum summertime soil moisture. Calculated over days when mean soil temperature is > 1°C in June-August | VWC% |
| moist_sd_summer | Moist | Standard deviation of summertime soil moisture. Calculated over days when mean soil temperature is > 1°C in June-August | VWC% |
| moist_cv_summer | Moist | Coefficient of variation of summertime soil moisture. Calculated over days when mean soil temperature is > 1°C in June-August | VWC% |
| moist_mean_july | Moist | Mean summertime soil moisture. Calculated over days when mean soil temperature is > 1°C in July | VWC% |
| moist_max_july | Moist | Maximum summertime soil moisture. Calculated over days when mean soil temperature is > 1°C in July | VWC% |
| moist_min_july | Moist | Minimum summertime soil moisture. Calculated over days when mean soil temperature is > 1°C in July | VWC% |
| moist_sd_july | Moist | Standard deviation of summertime soil moisture. Calculated over days when mean soil temperature is > 1°C in July | VWC% |
| moist_cv_july | Moist | Coefficient of variation of summertime soil moisture. Calculated over days when mean soil temperature is > 1°C in July | VWC% |

**Figure 2.**
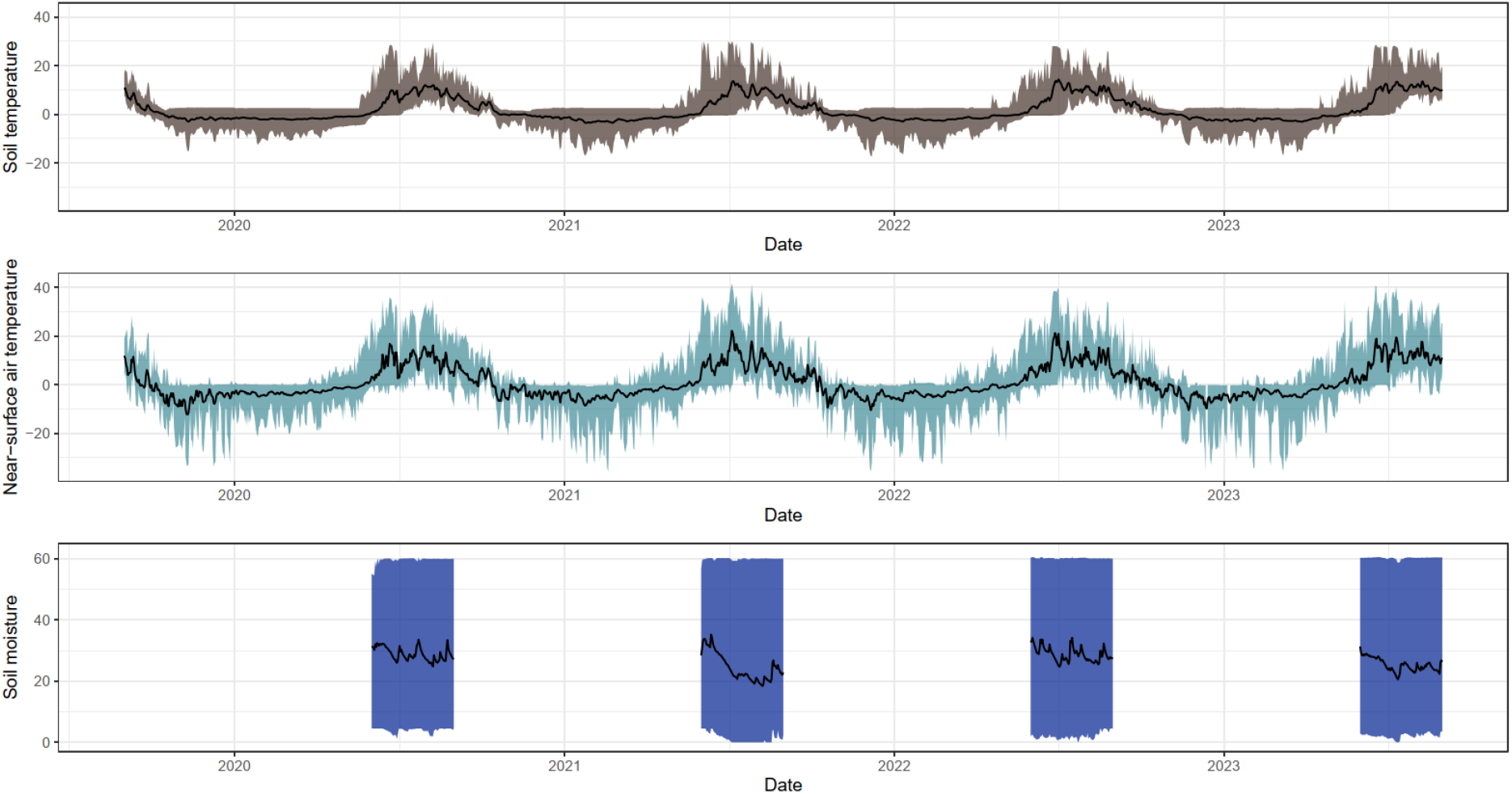
Variation in soil temperature, near surface air temperature and soil moisture over the study period from September 2019 to August 2023. Black lines represent daily averages and the coloured ribbons represent the range between the minimum and maximum values. Values were calculated from daily-aggregated imputed time series data across 430 loggers.

Loggers yielding less than one hydrological year of quality-filtered data (or one full summer in case of soil moisture) were removed from the remaining analyses, resulting in 430 loggers for air temperature variables, 428 for soil temperature variables and 422 for soil moisture variables. For these selected loggers, any missing observations (e.g., due to later installation date, malfunctioning, or a heaved-out logger) were imputed with a Random Forest model with functions from *MissForest* R package (Stekhoven and Buhlmann, 2012). Imputations were done separately for the daily means, maximums and minimums of T1 and T3 temperatures and for daily means of soil moisture, respectively. Imputation included only the logger data (for example, relating the daily mean T1 time series across all the loggers), and is thus completely independent from the environmental predictors used in the later modelling parts of the study. Performance of the imputations was evaluated with a leave-month-out cross-validation procedure, where we randomly selected a month for 20 random study sites which were withdrawn from the data used in the following imputation and then compared the withdrawn true values to the modelled (imputed) record. This procedure was repeated 10 times for each imputed variable (e.g., daily T1 means), so the evaluation was based on 200 withdrawn months for each imputed variable (see Table S1 for summary of the cross validation statistics).

### 2.3. Environmental predictors

We calculated a total of 72 spatially continuous variable candidates to be used as predictors for the microclimate response variables (Table 2). The predictors reflect various aspects of topography, snow conditions, vegetation structure and land cover types. We used the *terra* package (Hijmans, 2022) to work with raster data in R, *rsagacmd* package (Pawley, 2018) as an R interface for *SAGA-GIS* software (v8.5.1)(Conrad et al., 2015) and *whitebox* package (Wu and Brown, 2022) as an interface to *WhiteboxTools* software (v1.4.0)(Lindsay, 2016) to calculate the topographic and distance based variables. The *lidR* package (Roussel and Auty, 2019) was used to work with Light Detection and Ranging (LiDAR) data.

**Table 2.** Environmental predictors of the microclimate variables.

| Name | Data source | Explanation |
| --- | --- | --- |
| dem | LiDAR | Digital elevation model at 3-metre spatial resolution. Also used for calculating the rest of the topography-based variables. |
| pisr_1 ...<br>pisr_12 | LiDAR | Monthly potential incoming solar radiation assuming clear sky conditions. Sky view factor was calculated with a 1 km search radius. Calculated for the 15th day of each month with a temporal interval of one hour. |
| pisr20_1 ...<br>pisr20_12 | LiDAR | Monthly potential incoming solar radiation. Calculated as above, but smoothed (averaged) with a 20 m radius moving window. |
| tpi15 | LiDAR | Topographic position index with a search window of 15 m radius. The index is the difference of the elevation of the focal grid cell and the mean of the neighbouring cells. |
| tpi30 | LiDAR | Topographic position index with a search window of 30 m radius. |
| tpi90 | LiDAR | Topographic position index with a search window of 90 m radius. |
| tpi250 | LiDAR | Topographic position index with a search window of 250 m radius and by using a DEM aggregated to 15 m resolution. |
| tpi500 | LiDAR | Topographic position index with a search window of 500 m radius and by using a DEM aggregated to 15 m resolution. |
| tpi1000 | LiDAR | Topographic position index with a search window of 1000 m radius and by using a DEM aggregated to 15 m resolution. |
| slope | LiDAR | Slope steepness in degrees. |
| windexp | LiDAR | Wind exposition index describing the topographic openness with a 500-m radius search window. |
| windexp100 | LiDAR | Wind exposition index describing the topographic openness with a 100-m radius search window. |
| swi | LiDAR | Saga wetness index with default parameters (suction = 10, slope_type = catchment slope). |
| swi_suct2 | LiDAR | Saga wetness index with suction parameter set as 10 and slope type as local slope. |
| swi_suct8 | LiDAR | Saga wetness index with suction parameter set as 8 and slope type as local slope. |
| swi_suct64 | LiDAR | Saga wetness index with suction parameter set as 64 and slope type as local slope. |
| swi_suct256 | LiDAR | Saga wetness index with suction parameter set as 256 and slope type as local slope. |
| TWI_d8 | LiDAR | Topographic wetness index with a deterministic D8 flow-routing algorithm. |
| TWI_Dinf | LiDAR | Topographic wetness index with a multiple-flow-direction D-infinity flow-routing algorithm. |
| TWI_FD8f | LiDAR | Topographic wetness index with a multiple-flow-direction FD8 Freeman flow-routing algorithm. |
| disttowater | Water vectors | Euclidean distance to the closest waterbody (lakes, ponds, rivers and creeks derived from the topographic database of Finland). |
| watereffect | Water vectors | Slope-penalised distance to the closest waterbody. Slope included as cost surface when searching for the shortest cost distance to waterbodies. |
| scd | Satellite imagery | Snow cover duration, the average day of year when snow melts. |
| snowresi | Satellite imagery & LiDAR | The residuals of SCD ~ Elevation linear model, i.e., snow cover duration without the effect of elevation. |
| snoweffect | Satellite imagery & LiDAR | The uphill snow patch index, describes if a cell will likely be fed by meltwater from nearby uphill snow patches. |
| NDVI_max | Satellite imagery | Maximum normalised difference vegetation index, a proxy for vegetation greenness and biomass. |
| NDVI_median | Satellite imagery | Median NDVI during snow free season. |
| ndviresi | Satellite imagery & LiDAR | The residuals of NDVI_max ~ Elevation linear model, i.e., NDVI without the effect of elevation. |
| chm | LiDAR | Canopy height (cm). |
| ccv5m | LiDAR | Canopy cover (%) considering trees higher than 5 metres. |
| ccv2m | LiDAR | Canopy cover (%) considering trees higher than 2 metres. |
| shrubs | LiDAR | Shrub cover (%) considering vegetation between 0.25 and 1 metre, height typical for the shrubs in the study area. |
| Pseudo_albedo | Satellite imagery | Maximum surface reflectances summarised over the eight spectral bands in PlanetScope satellite data. |
| PCA1 ... PCA15 | Satellite imagery | First 15 principal components of spectral information in three PlanetScope satellite images. X = 1,2,3... 15. |

The topography predictors are based on a national digital elevation model (DEM) produced by the National Land Survey of Finland at 2-metre spatial resolution. The DEM is based on LiDAR data. The same LiDAR data was used to calculate predictors regarding vegetation structure and height. All used satellite data are multitemporal PlanetScope imagery (Planet Team, 2017). PlanetScope imagery has the largest pixel size (3 metres) of all material used to calculate the predictors, and thus, all other predictors were eventually resampled to the same 3-metre spatial resolution.

Monthly potential incoming solar radiation (pisr) was calculated with the Potential solar radiation tool in *SAGA-GIS* (Böhner and Antonić, 2009). We selected the 15^th^ day of each month and calculated the radiation for those dates with a one hour interval. The algorithm takes into account the surrounding topography of a focal pixel by including the Sky View Factor which we calculated with a 1-km search radius. All other parameters were kept as defaults.

Topographic position index (tpi) relates the elevation of the focal pixel to the average elevation within its neighbourhood with a defined search radius (Guisan et al., 1999). We used a range of search radii (Table 2), because tpis with different radii might reflect different physiographic processes. We used the *Topographic Position Index* tool in *SAGA-GIS* to calculate the indices.

We calculated multiple topographic wetness indices, which mimic the water flow and its accumulation across landscapes. These indices are commonly used as proxies for soil moisture conditions (Kopecky and Cizkova, 2010). Different wetness index algorithms result in differing spatial patterns in the index, which is why we included several wetness index options as predictor candidates. We used Saga Wetness Index (swi) algorithm (Böhner et al., 2002) with a range of suction parameters and also the original topographic wetness index (twi) algorithm (Beven and Kirkby, 1979) calculated with several flow routing algorithms (details are provided in Table 2; see Riihimäki et al. 2021 for more information about the wetness indices and their empirical test in Kilpisjärvi region). Before calculating the wetness indices, we filled possible sinks in the DEM with the *Fill Sinks XXL* tool in *SAGA-GIS* (Wang and Liu, 2006). Saga wetness indices were calculated with the *SAGA Wetness Index* tool in SAGA-GIS. Other wetness indices were calculated with the *Wetness Index* tool in *WhiteboxTools* software by preprocessing the inputted flow accumulation rasters with the *D8 flow accumulation*, *D-infinity flow accumulation* and *Fd8 flow accumulation* tools.

Slope is the local slope angle calculated with the *Slope, Aspect, Curvature* tool in *SAGA-GIS* by using the *9 parameter 2nd order polynomial* method (Zevenbergen and Thorne, 1987).

Wind exposition index describes the topographic openness around the focal grid cell with a specified search radius (Böhner and Antonić, 2009). We used the *Wind Exposition Index* tool in *SAGA-GIS* with two search radii: 100 and 500 metres.

Distance to water bodies and rivers were calculated with two methods: 1) a simple euclidean distance to the nearest water feature in the topographic database of Finland; and 2) a cost-distance to the water features where the local slope was used as a cost-surface because the effect of surface waters likely reach further on flat areas (see details of the method in Rissanen et al. 2023). The predictors were calculated by using the *Proximity Grid* and *Accumulated Cost* tools in *SAGA-GIS*.

Snow cover duration (*scd*) was constructed from 306 satellite images acquired in 2017–2021 (see Kemppinen & Niittynen (2022) for a detailed description of the method). In summary, all the individual multispectral PlanetScope satellite images were cleaned for cloud effects and binarised pixel by pixel to snow (or no-snow) maps. Then, we determined the pixel-wise average snow melting date by regressing the binary snow information to day-of-year values with a binomial generalised linear model. Furthermore, we calculated the residuals of a linear regression between elevation and scd to calculate a variable with information of snow cover duration independent of the altitudinal effect (predictor called *snowresi*). We also calculated an uphill distance from snowbed areas (*snoweffect*) to account for the impacts of melting water from the late-lying snow patches. Here, we first inverted the DEM and used the *Downslope distance to stream* tool in *WhiteboxTools* software to calculate the uphill distance to the closest snowbed cell. If the distance was greater than 100 metres, we set the distance value to 100. We iterated the calculation with a range of day-of-year thresholds (from 150 to 258 with 7-day intervals) to determine the snowbeds. Subsequently, we calculated a sum of all distance rasters produced along the iterations. Finally, we inverted the resulting raster so that higher values indicate the more likely influence of the nearby uphill snowbed.

We selected 19 completely or nearly cloud-free PlanetScope images from peak summer (from late June to early August) in 2017–2022 for deriving spectral information of the land surfaces. We removed cloudy pixels by using manually digitised cloud masks and we also removed pixels where the blue band reflectance was over 0.5 (i.e., unrealistically high for any bare soil, rock or vegetated surface). Next, we calculated a Normalised difference vegetation index (NDVI, a normalised ratio of red and near-infrared spectral bands) of each of the images, and finally, we determined the maximum and median NDVI values across all images for each pixel (predictors called *NDVI_max* and *NDVI_median*). Furthermore, we calculated the residuals of a linear regression between elevation and *NDVI_max* to calculate a variable where we have vegetation greenness information independent of the altitudinal effect (*ndviresi*).

We used the same 19 PlanetScope images as described above to calculate pseudo-albedo of the surfaces to characterise the brightness of the surfaces. Here, we used the same procedure as with NDVI analyses to mask clouds. We calculated a mean reflectance over all the eight spectral bands in the imagery for each image separately, and finally, we calculated the pixel-wise maximum mean reflectance across all 19 images.

We also conducted principal component analysis (PCA) for the PlanetScope images. Here, we selected three completely cloud-free images (acquired in 2022-07-02, 2022-07-30, and 2022-09-08) and masked all water surfaces from the images. Next, we stacked the three images together and calculated normalised difference indices between all possible pairs of bands within the image stack. We selected three of the resulting indices (Red ∼ NIR [i.e., NDVI], Green ∼ Red, and Yellow ∼ NIR) and calculated a sum of differences in these indices across the three imaging dates. Next, we stacked all the resulting indices and summed the difference rasters. This image stack with 87 layers would have too many pixels to run the PCA analyses, which is why we draw a random sample of 50,000 pixels. These pixels were used to run the PCA analyses. Finally, the resulting PCA model was used to predict the PCA values to the whole raster stack to produce continuous variables to be used as predictors in the microclimatic models. We selected the first 15 PCA components as predictor candidates. We used the normalised difference indices between the bands instead of the raw reflectance values in the PCA because the latter is more prone to topographical effects (i.e., shadows), and may be thus less sensitive to the actual differences in the land surface properties than the band ratios.

Vegetation height and canopy cover variables were generated from the Finnish national LiDAR data produced and preprocessed by the National Land Survey of Finland. The LiDAR data was read into R and the point-cloud elevation readings were normalised (i.e., the effect of the ground surface elevation removed with spatial interpolation based on a Delaunay triangulation). Next, the points higher than 25 metres were removed and the grid_canopy function in the *lidR* R package was finally used to calculate the vegetation height at an initial 1-m resolution. Vegetation height algorithms produced unrealistically high values on steep slopes. Thus, we set the vegetation height as zero if the slope angle was over 60 degrees, NDVI was under 0.4 (indicating sparse vegetation), or when simultaneously the slope was over 30 degrees and NDVI under 0.6. These thresholds were decided subjectively by simply visualising and exploring the data. Furthermore, the vegetation height was post-processed into three additional variables: 1) canopy cover of vegetation higher than 5 metres (i.e., the number of 1-m pixels with > 5m vegetation within a 6-metre radii circular search window); 2) same as above but considering vegetation over 2 metres; and 3) same as above but considering vegetation between 0.25 and 1 metres in height (i.e., the shrub cover).

### 2.4. Statistical analyses

We started the modelling with 72 predictors, but many of them showed strong collinearities. Having too many predictors may also lead to overfitting the models. Thus, we performed the following steps to prune the set of predictors.

1. We used the *nearZeroVar* function from *caret* package (Kuhn, 2021) to exclude predictors with minimal or zero variability. This step excluded potential solar radiation of January and December, i.e., the months with nonexistent solar radiation during the polar night.
2. Next we used the *findCorrelation* function from *caret* package to exclude bivariate correlations higher than |0.9|. If two predictors have a high absolute correlation, the function evaluates the mean absolute correlation of each predictor across all other predictors and removes the predictor with the higher mean absolute correlation. This excluded many of the monthly potential solar radiation predictions as well as several variants of the topographic wetness indices.
3. Finally, we conducted a Random Forest based recursive feature elimination procedure separately for each one of the response variables with the *rfeControl* function from *caret*. Here, the algorithm excludes predictors with a simple backwards selection to maximise the cross-validated (random 10-fold cross-validation repeated four times) predictive accuracy of the Random Forest model.

The remaining set of predictors was then used in the next step, in which we fitted the final predictive model. Here, we fitted Random Forest regression models separately for each response variable with the *ranger* R package (Wright and Ziegler, 2017) wrapped with the *train* function of *caret* package (Kuhn, 2021). To achieve the best-performing model we tuned the hyperparameters (mtry = 1,2,3,4,5,6,7 and 8; min.node.size = 1,2,3,4 and 5) with a random 10-fold cross-validation by minimising the RMSE. We recorded the cross-validation statistics of the best performing model and fitted a final full model (all data points included) with the tuned hyperparameters. This final model was then used to make predictions into the predictor raster stack to produce the continuous microclimatic surfaces for the whole study area at a 3-metre spatial resolution.

Many of the predictors used here are directly linked to the physical processes that regulate local temperatures and soil moisture regimes (Greiser et al., 2018; Kearney et al., 2020). Yet, many predictors are also more indirect and may only be correlated in space with the underlying mechanisms that control microclimate. We also selected a machine learning model instead of a more simple statistical modelling method which could be more straightforward to interpret. Both of these aspects might compromise, for example, the transferability and extrapolative capability of the models. However, here our main purpose was to build well-predicting models without any objective of extrapolation in space or time, nor any intent to explore the relationships between response and predictor variables more thoroughly. For purely predictive purposes, machine learning models have been shown to perform well (Bzdok et al., 2018).

## 3. Data records

The 63 microclimate rasters (see examples in Figure 3) are available to download in Zenodo repository (https://doi.org/10.5281/zenodo.10897219; Niittynen et al., 2024) as separate GeoTiff files, which is a geolocated raster image format. The naming of the files follows the nomenclature in Table 1. Also, all the predictor rasters are available as GeoTiffs in the same repository. The metadata of the microclimate and predictor rasters (effectively the same information as in Table 1 and 2) are available as two machine-readable text files. All code and data relevant to reproduce the results of this study are available in a Github repository (https://github.com/poniitty/Model_Kilpisjarvi_microclimates) and a stable version will be posted in the Zenodo data repository upon acceptance of the manuscript.

**Figure 3.**
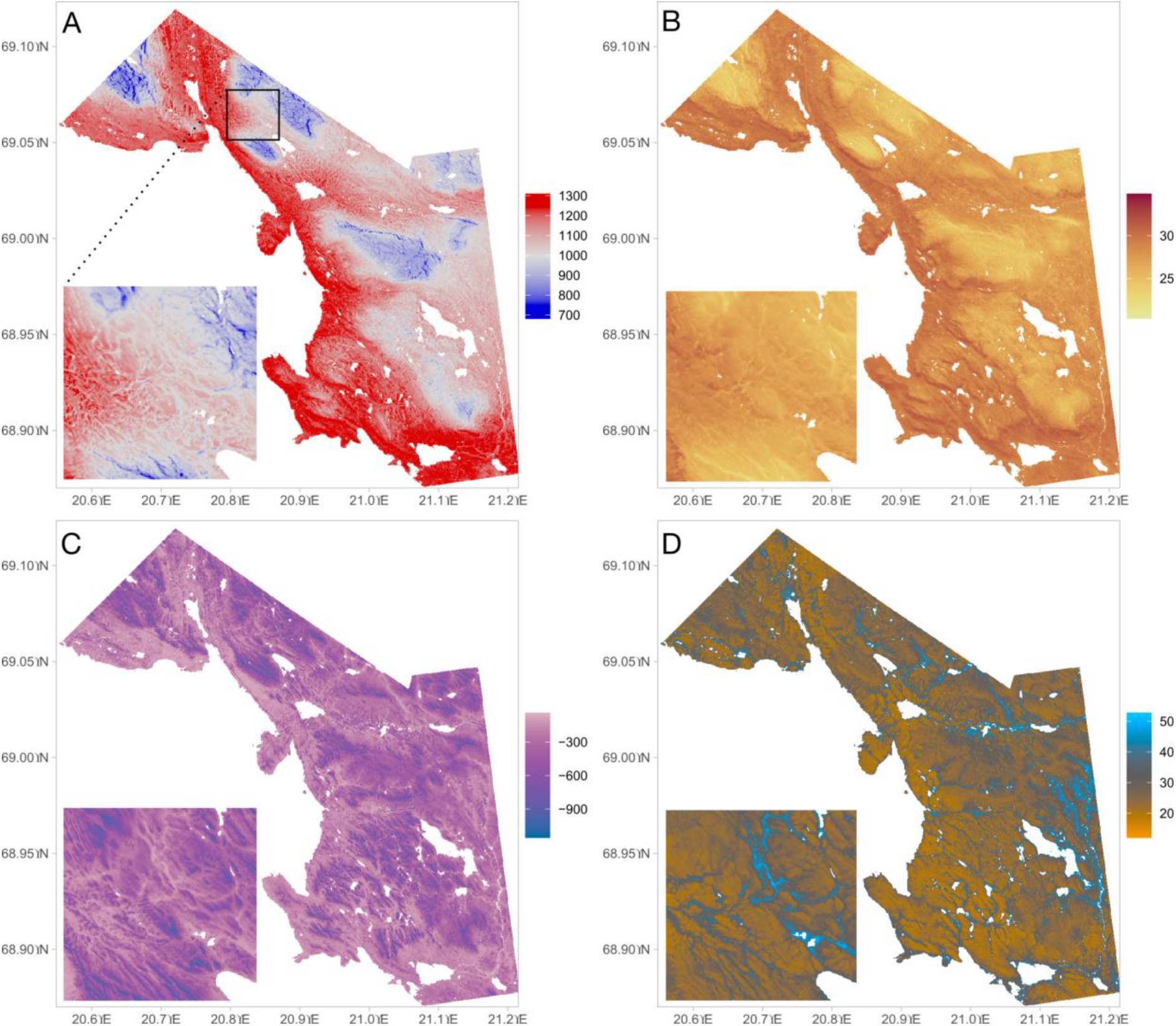
Predicted microclimate surfaces for four example variables. A) T3_GDD3, Near-surface air temperature growing degree days; B) T3_max, Maximum near-surface air temperature; C) T1_FDD, Soil temperature freezing degree days; D) moist_mean_summer, Summer-time mean soil moisture.

## 4. Technical validation

In general, the cross-validation statistics indicated that the models performed well, but also that there was large variability across the response variables (**Table 3**). On average, the cross-validated R^2^ value was 0.61 (median 0.64), which can be considered good when compared to previous studies in high-latitude regions with relatively similar study settings (Aalto et al., 2022; Greiser et al., 2018; Kemppinen et al., 2023). The best performing models had response variables which described summer and early autumn conditions.The worst-performing models were related to the freeze-thaw frequency in the soil and to temporal variability in soil moisture.

**Table 3.** Cross-validated model performance scores for the modelled microclimate variables. R^2^ = coefficient of determination; RMSE = Root mean squared error; MAE = Mean absolute error.

| Variable | $R^2$ | RMSE | MAE |
| --- | --- | --- | --- |
| T3_mean_9 | 0.936 | 0.21 | 0.16 |
| T3_mean_8 | 0.879 | 0.27 | 0.20 |
| T3_GDD5_days | 0.876 | 4.25 | 3.27 |
| T3_GDD3 | 0.868 | 55.29 | 42.08 |
| T3_TDD | 0.867 | 55.77 | 42.62 |
| T1_mean_9 | 0.864 | 0.40 | 0.31 |
| T3_GDD5 | 0.863 | 55.64 | 41.33 |
| T3_GDD3_days | 0.850 | 4.81 | 3.71 |
| T3_mean_7 | 0.839 | 0.36 | 0.25 |
| T1_GDD3_days | 0.798 | 6.19 | 4.83 |
| eos3 | 0.794 | 1.70 | 1.23 |
| T1_mean | 0.781 | 0.51 | 0.39 |
| T1_mean_4 | 0.773 | 0.58 | 0.44 |
| T3_mean_6 | 0.771 | 01.04 | 0.75 |
| T3_mean_5 | 0.769 | 0.48 | 0.37 |
| T3_mean_4 | 0.767 | 0.62 | 0.48 |
| T1_mean_10 | 0.755 | 0.42 | 0.33 |
| T1_mean_3 | 0.753 | 1.01 | 0.77 |
| T3_mean_3 | 0.723 | 1.21 | 0.93 |
| T1_mean_2 | 0.722 | 1.14 | 0.86 |
| T3_mean | 0.721 | 0.61 | 0.48 |
| T1_GDD5_days | 0.721 | 7.23 | 5.66 |
| T1_FDD | 0.713 | 172.82 | 130.91 |
| T3_max | 0.710 | 1.70 | 1.20 |
| T3_mean_10 | 0.707 | 0.43 | 0.32 |
| eos1 | 0.685 | 1.68 | 1.12 |
| T3_mean_2 | 0.677 | 1.50 | 1.17 |
| T1_mean_1 | 0.663 | 1.19 | 0.90 |
| T1_min | 0.654 | 1.80 | 1.41 |
| T1_mean_5 | 0.644 | 0.44 | 0.30 |
| T3_FDD | 0.643 | 235.07 | 185.09 |
| sos1 | 0.641 | 5.73 | 4.29 |
| T1_TDD | 0.637 | 105.17 | 80.85 |
| T1_GDD3 | 0.633 | 104.02 | 79.53 |
| T3_mean_1 | 0.621 | 1.65 | 1.31 |
| T1_mean_12 | 0.614 | 1.10 | 0.81 |
| T1_mean_8 | 0.607 | 0.69 | 0.52 |
| sos3 | 0.603 | 5.55 | 4.39 |
| T1_mean_11 | 0.596 | 0.70 | 0.52 |
| T1_GDD5 | 0.582 | 114.26 | 87.06 |
| T3_ft | 0.581 | 19.78 | 14.60 |
| T3_FDD_days | 0.550 | 9.90 | 7.38 |
| T3_TDD_days | 0.542 | 10.10 | 7.49 |
| T3_mean_12 | 0.515 | 1.60 | 1.27 |
| T1_mean_6 | 0.509 | 1.47 | 1.16 |
| T1_FDD_days | 0.495 | 23.33 | 14.48 |
| T3_frost | 0.456 | 4.75 | 3.49 |
| moist_mean_summer | 0.444 | 8.66 | 6.87 |
| T3_min | 0.441 | 3.20 | 2.52 |
| moist_mean_july | 0.440 | 8.78 | 6.95 |
| T1_TDD_days | 0.439 | 23.83 | 14.60 |
| moist_min_july | 0.431 | 8.73 | 6.81 |
| T3_mean_11 | 0.423 | 1.03 | 0.82 |
| moist_max_july | 0.420 | 9.10 | 7.33 |
| T1_max | 0.417 | 2.21 | 1.70 |
| moist_min_summer | 0.416 | 8.54 | 6.66 |
| T1_mean_7 | 0.410 | 1.08 | 0.83 |
| moist_max_summer | 0.401 | 10.13 | 8.37 |
| moist_sd_summer | 0.201 | 2.18 | 1.65 |
| moist_cv_summer | 0.175 | 0.09 | 0.07 |
| moist_cv_july | 0.150 | 0.06 | 0.05 |
| T1_ft | 0.149 | 4.86 | 3.22 |
| moist_sd_july | 0.072 | 1.28 | 0.97 |

Furthermore, the correlation structures remained similar across the pairs of microclimate variables when calculated either from the raw microclimate measurements or from the predicted microclimate rasters (**Figure S2**). The mean absolute difference between the correlation coefficients was 0.15 (median 0.13). This indicates that even though the model performance varies greatly, the relationships across different aspects of microclimate do not get strongly biassed in the modelling and projecting steps. The largest changes in the pairwise correlations were in general associated with the variables with the lowest cross-validated R^2^.

## 5. Usage notes

To make optimal use of the dataset, the user should first consider which variables (out of the many calculated) are most relevant for their specific study questions (**Table 1**). Next, the user should evaluate how well those variables performed in the technical validation of this study, with a preference for variables associated with good model performance (**Table 3**). Finally, the user should consider the correlation structure of the variables, aiming to exclude highly correlated variables (Figure 4). For example, out of the 63 candidate variables, it is possible to select seven variables (moist_cv_summer, moist_mean_summer, moist_sd_summer, sos1, T1_max, T3_mean_9, and T3_min) with no pairwise correlations exceeding |0.63|. Nonetheless, some of these variables showed poor model performance, of which the user should be aware of.

**Figure 4.**
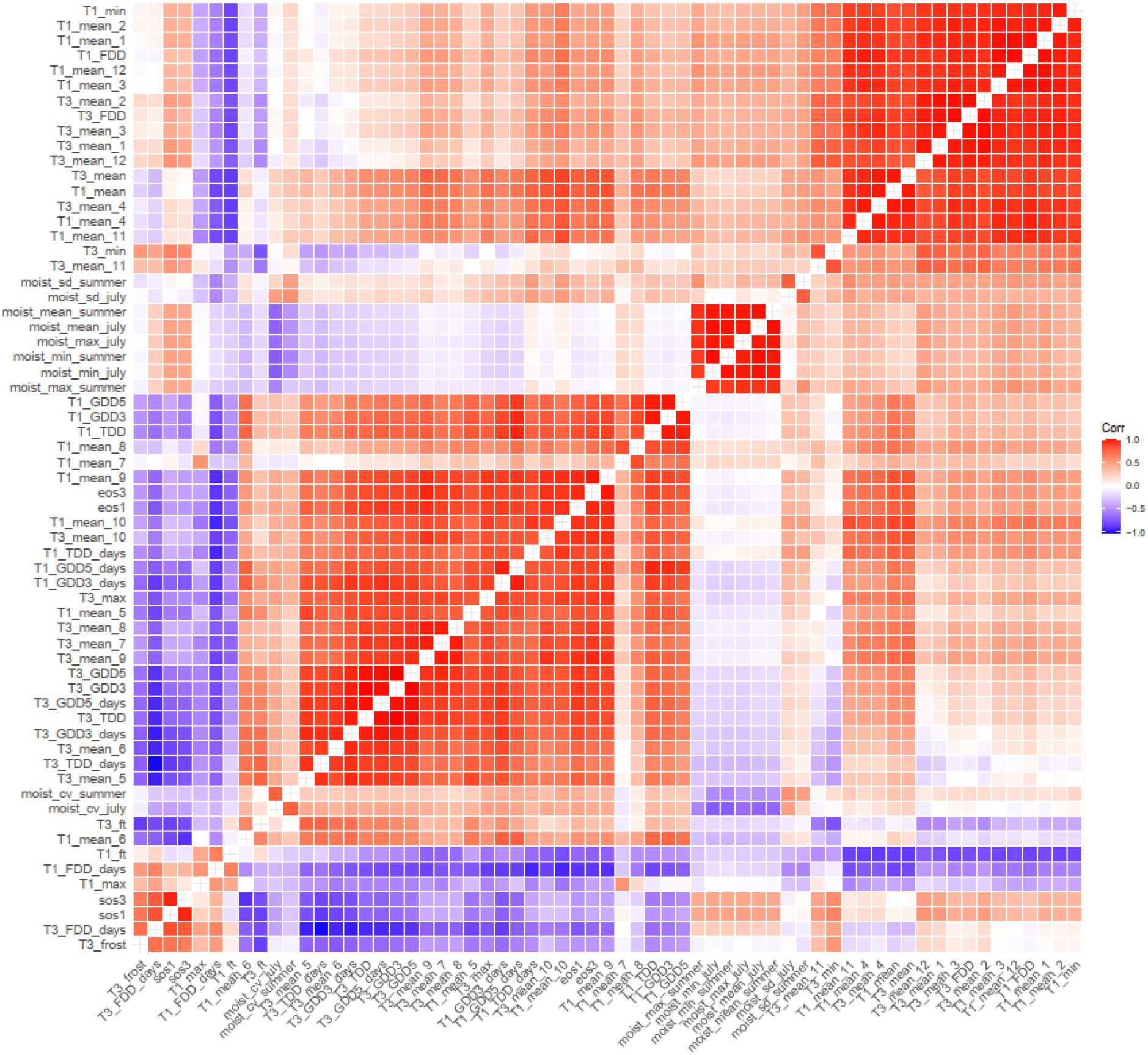
A correlation matrix across all modelled microclimate variables. The correlation coefficients were calculated from a random sample of 10,000 pixels extracted from the projected gridded microclimate surfaces.

As the model calibration data comprehensively covers the main environmental gradients within the study area, extrapolation beyond the data range will generally not be needed. In fact, models based on decision trees cannot be extrapolated at all. However, one environmental gradient was suboptimally covered by the studied sites: the very late-melting snowbeds (e.g., habitats from which snow melts in August) were not present in the microclimate data. As a result, the spatial predictions will likely overestimate the length of summer (and corresponding summer-time variables) in those locations. Thus, any user working on very late-melting habitats should be aware of this issue. Additionally, there can be some very specific local microclimate pockets, such as springs and their immediate surroundings, which are neither represented in the logger data nor in the predictors.

## Acknowledgements

We thank the staff of Kilpisjärvi research station, fieldwork assistants, and the Finnish IT Center for Science (CSC). PN thanks the Research Council of Finland (decision no. 347558), the Nessling Foundation and the Finnish Cultural Foundation for making this study possible. JK acknowledges the Research Council of Finland (349606), Maa-ja vesitekniikan tuki ry., Tiina and Antti Herlin Foundation, Nordenskiöld-samfundet, and the University of Oulu (Arctic Interactions, Arctic Researchers’ Network, and Biodiverse Anthropocenes). TR was funded by the European Research Council (ERC) under the European Union’s Horizon 2020 research and innovation programme (ERC-synergy grant 856506—LIFEPLAN). TR and PDLPA were further supported by a career support grant (to TR) from the Vice Chancellor of SLU, by a grant from the research council FORMAS (grant FR-2019/0007, to TR) and by a grant from the Academy of Finland (grant 322266, to TR). JA and ML were funded by the Academy of Finland (grant 342890). We are grateful for Mitro Müller, Sakari Sarjakoski and Heli Kainulainen who helped setting out and maintaining the study design 1. TM was funded by Finnish Cultural Foundation and Biodiverse Anthropocenes project (336449) supported by The University of Oulu and The Research Council of Finland Profi6 funding. JAhuhta and HT were funded by The Research Council of Finland (grant 322652). We are grateful to Bastien Parisy, Helena Wirta, Mikko Tiusanen, Laerke Stewart and Tuomas Kankaanpää who helped in setting out and/or retrieving the loggers of study design 5.

## Supplementary materials

**Supplementary Figure S1.**
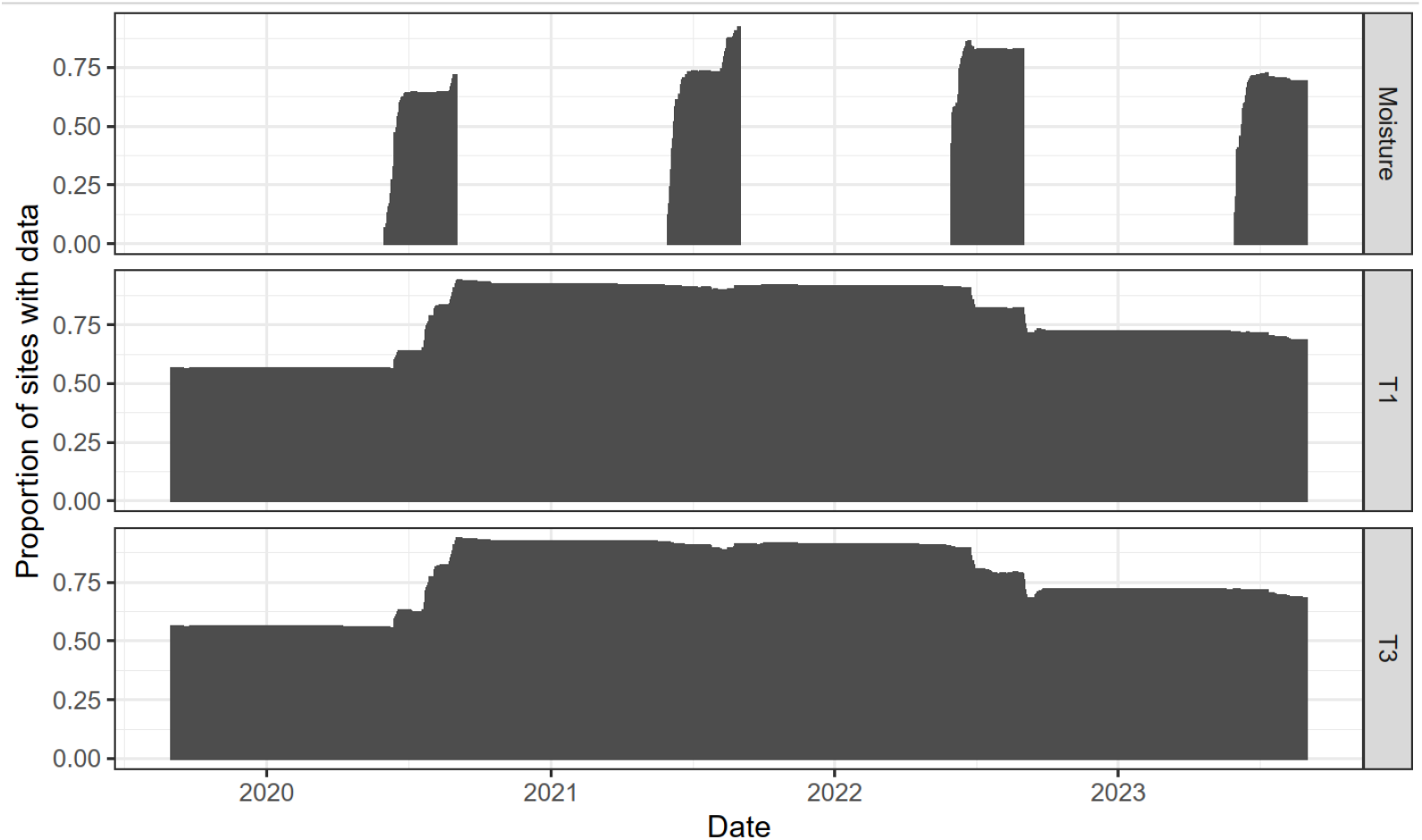
Proportion of the loggers with quality-filtered data on each day. T1 = soil temperature. T3 = near-surface air temperature. Moisture = soil moisture.

**Supplementary Table S1.** Cross-validated statistics of the imputation. Unit in temperature variables is ℃. Unit in soil moisture is volumetric water content (%). MAE = mean absolute error; MedAE = median absolute error; Bias = mean error; RMSE = root mean squared error

| Variable | MAE | MedAE | Bias | RMSE |
| --- | --- | --- | --- | --- |
| T1_max | 0.356 | 0.179 | 0.010 | 0.701 |
| T1_mean | 0.264 | 0.141 | -0.035 | 0.475 |
| T1_min | 0.290 | 0.149 | 0.022 | 0.524 |
| T3_max | 0.733 | 0.404 | -0.024 | 1.314 |
| T3_mean | 0.531 | 0.270 | -0.031 | 0.981 |
| T3_min | 0.575 | 0.330 | 0.052 | 0.952 |
| moist | 1.849 | 1.031 | 0.114 | 3.107 |

**Supplementary Figure S2.**
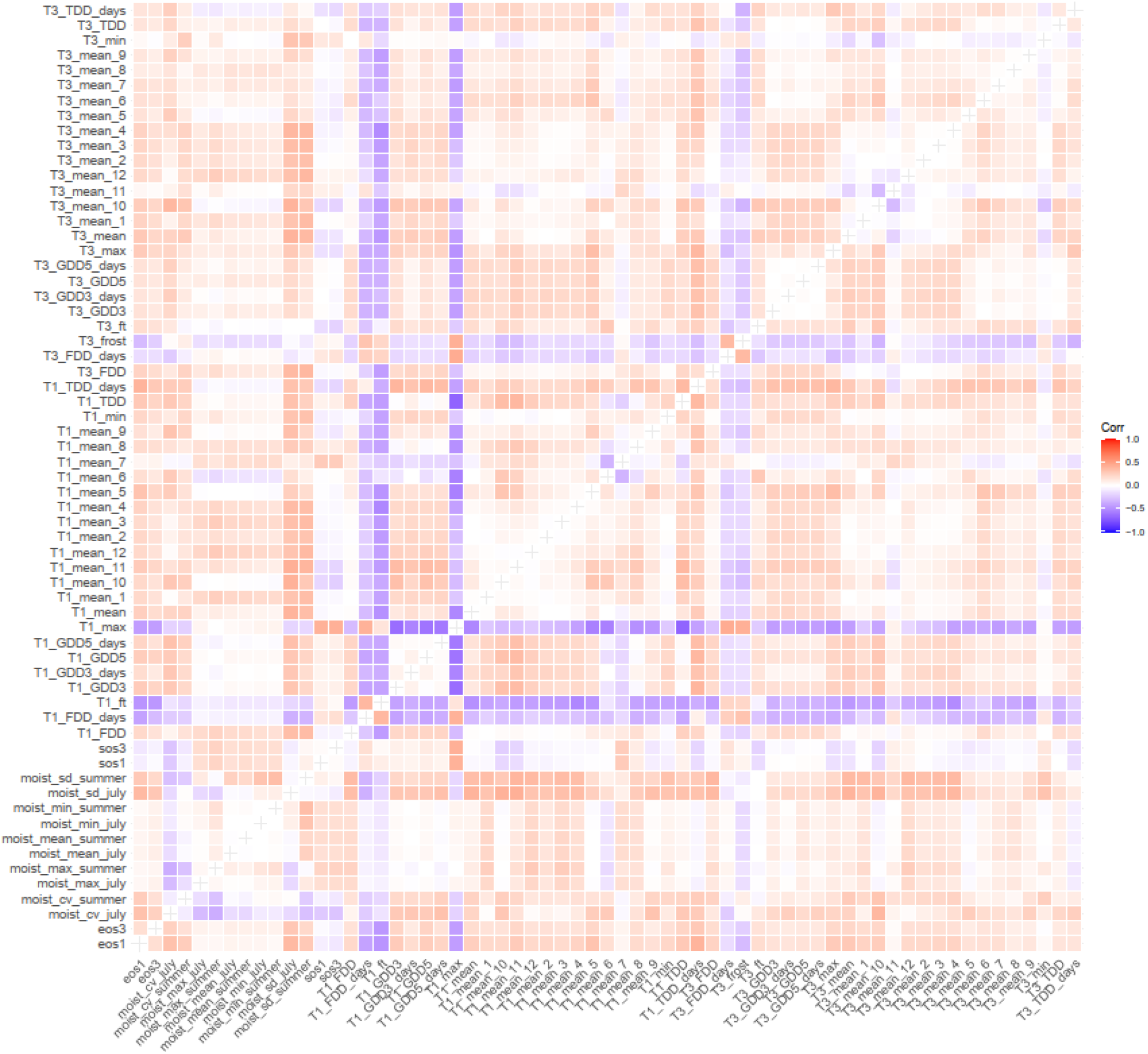
Difference in correlation coefficients calculated from the imputed logger data versus the modelled microclimate surfaces. All abbreviations and response variables are explained in detail in *Table 1*.

## References

1. Aalto, J., Luoto, M., 2014. Integrating climate and local factors for geomorphological distribution models. Earth Surf. Process. Landf. 39, 1729–1740. 10.1002/esp.3554

2. Aalto, J., Tyystjärvi, V., Niittynen, P., Kemppinen, J., Rissanen, T., Gregow, H., Luoto, M., 2022. Microclimate temperature variations from boreal forests to the tundra. Agric. For. Meteorol. 323, 109037. 10.1016/j.agrformet.2022.109037

3. Aho, J., Kalela, O., 1966. The spring migration of 1961 in the Norwegian lemming, Lemmus lemmus (L.), at Kilpisjärvi, Finnish Lapland. Ann. Zool. Fenn. 3, 53–65.

4. Beven, K.J., Kirkby, M.J., 1979. A physically based, variable contributing area model of basin hydrology. Hydrol. Sci. Bull. 24, 43–69. 10.1080/02626667909491834

5. Böhner, J., Antonić, O., 2009. Land-Surface Parameters Specific to Topo-Climatology, in: Hengl, T., Reuter, H.I. (Eds.), Developments in Soil Science, Geomorphometry. Elsevier, pp. 195–226. 10.1016/S0166-2481(08)00008-1

6. Böhner, J., Koethe, R., Conrad, O., Gross, J., Ringeler, A., Selige, T., 2002. Soil regionalisation by means of terrain analysis and process parameterisation. Eur. SOIL Bur. - Res. Rep. NO 7 213–222.

7. Bokhorst, S., Phoenix, G.K., Berg, M.P., Callaghan, T.V., Kirby-Lambert, C., Bjerke, J.W., 2015. Climatic and biotic extreme events moderate long-term responses of above- and belowground sub-Arctic heathland communities to climate change. Glob. Change Biol. 21, 4063–4075. 10.1111/gcb.13007

8. Bokhorst, S.F., Bjerke, J.W., Tommervik, H., Callaghan, T.V., Phoenix, G.K., 2009. Winter warming events damage sub-Arctic vegetation: consistent evidence from an experimental manipulation and a natural event. J. Ecol. 97, 1408–1415. 10.1111/j.1365-2745.2009.01554.x

9. Bramer, I., Anderson, B.J., Bennie, J., Bladon, A.J., De Frenne, P., Hemming, D., Hill, R.A., Kearney, M.R., Körner, C., Korstjens, A.H., Lenoir, J., Maclean, I.M.D., Marsh, C.D., Morecroft, M.D., Ohlemüller, R., Slater, H.D., Suggitt, A.J., Zellweger, F., Gillingham, P.K., 2018. Chapter Three - Advances in Monitoring and Modelling Climate at Ecologically Relevant Scales, in: Bohan, D.A., Dumbrell, A.J., Woodward, G., Jackson, M. (Eds.), Advances in Ecological Research, Next Generation Biomonitoring: Part 1. Academic Press, pp. 101–161. 10.1016/bs.aecr.2017.12.005

10. Burt, T.P., Butcher, D.P., 1985. Topographic Controls of Soil-Moisture Distributions. J. Soil Sci. 36, 469–486.

11. Bzdok, D., Altman, N., Krzywinski, M., 2018. Statistics versus machine learning. Nat. Methods 15, 233–234. 10.1038/nmeth.4642

12. Chen, J., Saunders, S.C., Crow, T.R., Naiman, R.J., Brosofske, K.D., Mroz, G.D., Brookshire, B.L., Franklin, J.F., 1999. Microclimate in Forest Ecosystem and Landscape Ecology: Variations in local climate can be used to monitor and compare the effects of different management regimes. BioScience 49, 288–297. 10.2307/1313612

13. Conrad, O., Bechtel, B., Dietrich, H., Fischer, E., Gerlitz, L., Wehberg, J., Wichmann, V., Böhner, J., 2015. System for Automated Geoscientific Analyses (SAGA) v. 2.1.4. Geosci. Model Dev. 8, 1991–2007. 10.5194/gmd-8-1991-2015

14. De Frenne, P., Lenoir, J., Luoto, M., Scheffers, B.R., Zellweger, F., Aalto, J., Ashcroft, M.B., Christiansen, D.M., Decocq, G., De Pauw, K., Govaert, S., Greiser, C., Gril, E., Hampe, A., Jucker, T., Klinges, D.H., Koelemeijer, I.A., Lembrechts, J.J., Marrec, R., Meeussen, C., Ogee, J.M., Tyystjarvi, V., Vangansbeke, P., Hylander, K., 2021. Forest microclimates and climate change: Importance, drivers and future research agenda. Glob. Change Biol. 27, 2279–2297. 10.1111/gcb.15569

15. Geiger, R., 1965. The climate near the ground, Cambridge: Harvard University Press. Cambridge: Harvard University Press.

16. Greiser, C., Meineri, E., Luoto, M., Ehrlen, J., Hylander, K., 2018. Monthly microclimate models in a managed boreal forest landscape. Agric. For. Meteorol. 250, 147–158. 10.1016/j.agrformet.2017.12.252

17. Guisan, A., Weiss, S.B., Weiss, A.D., 1999. GLM versus CCA spatial modeling of plant species distribution. Plant Ecol. 143, 107–122. 10.1023/a:1009841519580

18. Haapasaari, M., 1988. Oligotrophic heath vegetation of northern Fennoscandia and its zonation. Acta Bot. Fenn. 135.

19. Hannah, L., Flint, L., Syphard, A.D., Moritz, M.A., Buckley, L.B., McCullough, I.M., 2014. Fine-grain modeling of species’ response to climate change: holdouts, stepping-stones, and microrefugia. Trends Ecol. Evol. 29, 390–397. 10.1016/j.tree.2014.04.006

20. Hijmans, R.J., 2022. terra: Spatial Data Analysis.

21. Jokinen, P., Pirinen, P., Kaukoranta, J., Kangas, A., Alenius, P., Eriksson, P., Johansson, M., Wilkman, S., 2021. Climatological and oceanographic statistics of Finland 1991– 2020. Finn. Meteorol. Inst.

22. Kauhanen, H.O., 2013. Mountains of Kilpisjärvi host an abundance of threatened plants in Finnish Lapland. J Bot. Pacifica J. Plant Sci. 2, 43–52.

23. Kauppila, T., Salonen, V.-P., 1997. The effect of Holocene treeline fluctuations on the sediment chemistry of Lake Kilpisjärvi, Finland. J. Paleolimnol. 18, 145–163. 10.1023/A:1007978318562

24. Kearney, M.R., Gillingham, P.K., Bramer, I., Duffy, J.P., Maclean, I.M.D., 2020. A method for computing hourly, historical, terrain-corrected microclimate anywhere on earth. Methods Ecol. Evol. 11, 38–43. 10.1111/2041-210x.13330

25. Kemppinen, J., Niittynen, P., 2022. Microclimate relationships of intraspecific trait variation in sub-Arctic plants. Oikos 2022, e09507. 10.1111/oik.09507

26. Kemppinen, J., Niittynen, P., le Roux, P.C., Momberg, M., Happonen, K., Aalto, J., Rautakoski, H., Enquist, B.J., Vandvik, V., Halbritter, A.H., Maitner, B., Luoto, M., 2021a. Consistent trait-environment relationships within and across tundra plant communities. Nat. Ecol. Evol. 5, 458–467. 10.1038/s41559-021-01396-1

27. Kemppinen, J., Niittynen, P., Riihimaki, H., Luoto, M., 2018. Modelling soil moisture in a high-latitude landscape using LiDAR and soil data. Earth Surf. Process. Landf. 43, 1019–1031. 10.1002/esp.4301

28. Kemppinen, J., Niittynen, P., Rissanen, T., Tyystjärvi, V., Aalto, J., Luoto, M., 2023. Soil Moisture Variations From Boreal Forests to the Tundra. Water Resour. Res. 59, e2022WR032719. 10.1029/2022WR032719

29. Kemppinen, J., Niittynen, P., Virkkala, A.M., Happonen, K., Riihimaki, H., Aalto, J., Luoto, M., 2021b. Dwarf Shrubs Impact Tundra Soils: Drier, Colder, and Less Organic Carbon. Ecosystems In press. 10.1007/s10021-020-00589-2

30. King, L., Seppälä, M., 1987. Permafrost thickness and distribution in Finnish Lapland - results of geoelectrical soundings. Polarforsch. - 127–147.

31. Kopecky, M., Cizkova, S., 2010. Using topographic wetness index in vegetation ecology: does the algorithm matter? Appl. Veg. Sci. 13, 450–459. 10.1111/j.1654-109X.2010.01083.x

32. Kropp, H., Loranty, M.M., Natali, S., Kholodov, A.L., Abbott, B.W., Abermann, J., Blanc-Betes, E., Blok, D., Blume-Werry, G., Boike, J., Cahoon, C.M.P., Christiansen, C.T., Euskirchen, E.S., Frost, G.V., Goeckede, M., Gough, L., Hjorth, J., Hoje, T.T., Jones, B.M., Jorgenson, T., Juszak, I., Kokelj, S., Lund, M., Lafleur, P., Mamet, S., Mauritz, M., Michelsen, A., Myers-Smith, I.H., O’Donnell, J., Olefeldt, D., Phoenix, G.K., Rocha, A.V., Sannel, B., Schaepman-Strub, G., Sonnentag, O., Tape, K.D., Torn, M.S., Smith Vaughn, L., Sorensen, M., Williams, M., Wilson, C.J., 2016. Impacts of Vegetation on the Decoupling between Air and Soil Temperatures across the Arctic, in: EPIC3AGU Fall Meeting, San Francisco, 2016-12-12-2016-12-16AGU. Presented at the AGU Fall Meeting, AGU, San Francisco.

33. Kuhn, M., 2021. caret: Classification and Regression Training. R package version 6.0-89.

34. le Roux, P.C., Aalto, J., Luoto, M., 2013. Soil moisture’s underestimated role in climate change impact modelling in low-energy systems. Glob. Change Biol. 19, 2965–75. 10.1111/gcb.12286

35. Leppäranta, M., Lindgren, E., Shirasawa, K., 2016. The heat budget of Lake Kilpisjärvi in the Arctic tundra. Hydrol. Res. 48, 969–980. 10.2166/nh.2016.171

36. Lindsay, J.B., 2016. Whitebox GAT: A case study in geomorphometric analysis. Comput. Geosci. 95, 75–84.

37. Maclean, I.M.D., Duffy, J.P., Haesen, S., Govaert, S., De Frenne, P., Vanneste, T., Lenoir, J., Lembrechts, J.J., Rhodes, M.W., Van Meerbeek, K., 2021. On the measurement of microclimate. Methods Ecol. Evol. 12, 1397–1410. 10.1111/2041-210X.13627

38. Maliniemi, T., Kapfer, J., Saccone, P., Skog, A., Virtanen, R., 2018. Long-term vegetation changes of treeless heath communities in northern Fennoscandia: Links to climate change trends and reindeer grazing. J. Veg. Sci. Accepted Author Manuscript. 10.1111/jvs.12630

39. Niittynen, P., Heikkinen, R.K., Aalto, J., Guisan, A., Kemppinen, J., Luoto, M., 2020. Fine-scale tundra vegetation patterns are strongly related to winter thermal conditions. Nat. Clim. Change 10, 1143–U134. 10.1038/s41558-020-00916-4

40. Niittynen, P., Salminen, H., Peña-Aguilera, P., Aalto, J., Alahuhta, J., Luoto, M., Maliniemi, T., Rissanen, T., Roslin, T., Tukiainen, H., Tyystjärvi, V., Kemppinen, J., 2024. A Dataset: Gridded Microclimate Dataset from a Sub-Arctic Biodiversity Hotspot in Finland. 10.5281/zenodo.10897219

41. Opedal, O.H., Armbruster, W.S., Graae, B.J., 2015. Linking small-scale topography with microclimate, plant species diversity and intra-specific trait variation in an alpine landscape. Plant Ecol. Divers. 8, 305–315. 10.1080/17550874.2014.987330

42. Parmesan, C., Root, T.L., Willig, M.R., 2000. Impacts of extreme weather and climate on terrestrial biota. Bull. Am. Meteorol. Soc. 81, 443–450. 10.1175/1520-0477(2000)0810443:IOEWAC2.3.CO;2

43. Pawley, S., 2018. Rsagacmd: Linking R with the Open-Source “SAGA-GIS” Software. R package version 0.2.0.

44. Peña-Aguilera, P., Schmidt, N.M., Stewart, L., Parisy, B., van der Wal, R., Lindman, L., Vesterinen, E.J., Maclean, I.M.D., Kankaanpää, T., Wirta, H., Roslin, T., 2023. Consistent imprints of elevation, soil temperature and moisture on plant and arthropod communities across two subarctic landscapes. Insect Conserv. Divers. 16, 684–700. 10.1111/icad.12667

45. Pepin, N.C., Norris, J.R., 2005. An examination of the differences between surface and free-air temperature trend at high-elevation sites: Relationships with cloud cover, snow cover, and wind. J. Geophys. Res.-Atmospheres 110, D24112–D24112. 10.1029/2005JD006150

46. Planet Team, 2017. Planet Application Program Interface: In Space for Life on Earth [WWW Document]. URL https://api.planet.com

47. Potter, K.A., Woods, H.A., Pincebourde, S., 2013. Microclimatic challenges in global change biology. Glob. Change Biol. 19, 2932–2939. 10.1111/gcb.12257

48. R Core Team, 2023. R: A language and environment for statistical computing. R Foundation for Statistical Computing. Vienna Austria https://www.R-project.org/.

49. Raidla, V., Kaup, E., Ivask, J., 2015. Factors affecting the chemical composition of snowpack in the Kilpisjärvi area of North Scandinavia. Atmos. Environ. 118, 211–218. 10.1016/j.atmosenv.2015.07.043

50. Riihimäki, H., Kemppinen, J., Kopecký, M., Luoto, M., 2021. Topographic Wetness Index as a Proxy for Soil Moisture: The Importance of Flow-Routing Algorithm and Grid Resolution. Water Resour. Res. 57, e2021WR029871. 10.1029/2021WR029871

51. Rissanen, T., Aalto, A., Kainulainen, H., Kauppi, O., Niittynen, P., Soininen, J., Luoto, M., 2023. Local snow and fluvial conditions drive taxonomic, functional and phylogenetic plant diversity in tundra. Oikos 2023, e09998. 10.1111/oik.09998

52. Rixen, C., Høye, T.T., Macek, P., Aerts, R., Alatalo, J.M., Anderson, J.T., Arnold, P.A., Barrio, I.C., Bjerke, J.W., Björkman, M.P., Blok, D., Blume-Werry, G., Boike, J., Bokhorst, S., Carbognani, M., Christiansen, C.T., Convey, P., Cooper, E.J., Cornelissen, J.H.C., Coulson, S.J., Dorrepaal, E., Elberling, B., Elmendorf, S.C., Elphinstone, C., Forte, T.G.W., Frei, E.R., Geange, S.R., Gehrmann, F., Gibson, C., Grogan, P., Halbritter, A.H., Harte, J., Henry, G.H.R., Inouye, D.W., Irwin, R.E., Jespersen, G., Jónsdóttir, I.S., Jung, J.Y., Klinges, D.H., Kudo, G., Lämsä, J., Lee, H., Lembrechts, J.J., Lett, S., Lynn, J.S., Mann, H.M.R., Mastepanov, M., Morse, J., Myers-Smith, I.H., Olofsson, J., Paavola, R., Petraglia, A., Phoenix, G.K., Semenchuk, P., Siewert, M.B., Slatyer, R., Spasojevic, M.J., Suding, K., Sullivan, P., Thompson, K.L., Väisänen, M., Vandvik, V., Venn, S., Walz, J., Way, R., Welker, J.M., Wipf, S., Zong, S., 2022. Winters are changing: snow effects on Arctic and alpine tundra ecosystems. Arct. Sci. 8, 572–608. 10.1139/as-2020-0058

53. Roussel, J.-R., Auty, D., 2019. lidR: Airborne LiDAR Data Manipulation and Visualization for Forestry Applications.

54. Salminen, H., Tukiainen, H., Alahuhta, J., Hjort, J., Huusko, K., Grytnes, J.-A., Pacheco-Riaño, L.C., Kapfer, J., Virtanen, R., Maliniemi, T., 2023. Assessing the relation between geodiversity and species richness in mountain heaths and tundra landscapes. Landsc. Ecol. 38, 2227–2240. 10.1007/s10980-023-01702-1

55. Stekhoven, D.J., Buhlmann, P., 2012. MissForest-non-parametric missing value imputation for mixed-type data. Bioinformatics 28, 112–118. 10.1093/bioinformatics/btr597

56. Tyystjarvi, V., Kemppinen, J., Luoto, M., Aalto, T., Markkanen, T., Launiainen, S., Kieloaho, A.J., Aalto, J., 2022. Modelling spatio-temporal soil moisture dynamics in mountain tundra. Hydrol. Process. 36. 10.1002/hyp.14450

57. Wang, L., Liu, H., 2006. An efficient method for identifying and filling surface depressions in digital elevation models for hydrologic analysis and modelling. Int. J. Geogr. Inf. Sci. 20, 193–213. 10.1080/13658810500433453

58. Westermann, S., Østby, T.I., Gisnås, K., Schuler, T.V., Etzelmüller, B., 2015. A ground temperature map of the North Atlantic permafrost region based on remote sensing and reanalysis data. The Cryosphere 9, 1303–1319. 10.5194/tc-9-1303-2015

59. Wild, J., Kopecký, M., Macek, M., Šanda, M., Jankovec, J., Haase, T., 2019. Climate at ecologically relevant scales: A new temperature and soil moisture logger for long-term microclimate measurement. Agric. For. Meteorol. 268, 40–47. 10.1016/j.agrformet.2018.12.018

60. Winkler, M., Lamprecht, A., Steinbauer, K., Hulber, K., Theurillat, J.P., Breiner, F., Choler, P., Ertl, S., Giron, A.G., Rossi, G., Vittoz, P., Akhalkatsi, M., Bay, C., Alonso, J.L.B., Bergstrom, T., Carranza, M.L., Corcket, E., Dick, J., Erschbamer, B., Calzado, R.F., Fosaa, A.M., Gavilan, R.G., Ghosn, D., Gigauri, K., Huber, D., Kanka, R., Kazakis, G., Klipp, M., Kollar, J., Kudernatsch, T., Larsson, P., Mallaun, M., Michelsen, O., Moiseev, P., Moiseev, D., Molau, U., Mesa, J.M., di Cella, U.M., Nagy, L., Petey, M., Puscas, M., Rixen, C., Stanisci, A., Suen, M., Syverhuset, A.O., Tomaselli, M., Unterluggauer, P., Ursu, T., Villar, L., Gottfried, M., Pauli, H., 2016. The rich sides of mountain summits - a pan-European view on aspect preferences of alpine plants. J. Biogeogr. 43, 2261–2273. 10.1111/jbi.12835

61. Wright, M.N., Ziegler, A., 2017. ranger: A Fast Implementation of Random Forests for High Dimensional Data in C++ and R. J. Stat. Softw. 77, 1–17. 10.18637/jss.v077.i01

62. Wu, Q., Brown, A., 2022. “whitebox”: “WhiteboxTools” R Frontend.

63. Zevenbergen, L.W., Thorne, C.R., 1987. Quantitative analysis of land surface topography. Earth Surf. Process. Landf. 12, 47–56. 10.1002/esp.3290120107

64. Zhang, T.J., 2005. Influence of the seasonal snow cover on the ground thermal regime: An overview. Rev. Geophys. 43, RG4002. 10.1029/2004rg000157

